# BOUNTI: Brain vOlumetry and aUtomated parcellatioN for 3D feTal MRI

**DOI:** 10.1101/2023.04.18.537347

**Authors:** Alena U. Uus, Vanessa Kyriakopoulou, Antonios Makropoulos, Abi Fukami-Gartner, Daniel Cromb, Alice Davidson, Lucilio Cordero-Grande, Anthony N. Price, Irina Grigorescu, Logan Z. J. Williams, Emma C. Robinson, David Lloyd, Kuberan Pushparajah, Lisa Story, Jana Hutter, Serena J. Counsell, A. David Edwards, Mary A. Rutherford, Joseph V. Hajnal, Maria Deprez

**Author notes:** Email address:* (Alena U. Uus). Equal contribution (Alena U. Uus, Vanessa Kyriakopoulou).

## Abstract

Fetal MRI is widely used for quantitative brain volumetry studies. However, currently, there is a lack of universally accepted protocols for fetal brain parcellation and segmentation. Published clinical studies tend to use different segmentation approaches that also reportedly require significant amounts of time-consuming manual refinement. In this work, we propose to address this challenge by developing a new robust deep learning-based fetal brain segmentation pipeline for 3D T2w motion corrected brain images. At first, we defined a new refined brain tissue parcellation protocol with 19 regions-of-interest using the new fetal brain MRI atlas from the Developing Human Connectome Project. This protocol design was based on evidence from histological brain atlases, clear visibility of the structures in individual subject 3D T2w images and the clinical relevance to quantitative studies. It was then used as a basis for developing an automated deep learning brain tissue parcellation pipeline trained on 360 fetal MRI datasets with different acquisition parameters using semi-supervised approach with manually refined labels propagated from the atlas. The pipeline demonstrated robust performance for different acquisition protocols and GA ranges. Analysis of tissue volumetry for 390 normal participants (21-38 weeks gestational age range), scanned with three different acquisition protocols, did not reveal significant differences for major structures in the growth charts. Only minor errors were present in < 15% of cases thus significantly reducing the need for manual refinement. In addition, quantitative comparison between 65 fetuses with ventriculomegaly and 60 normal control cases were in agreement with the findings reported in our earlier work based on manual segmentations. These preliminary results support the feasibility of the proposed atlas-based deep learning approach for large-scale volumetric analysis. The created fetal brain volumetry centiles and a docker with the proposed pipeline are publicly available online at https://hub.docker.com/r/fetalsvrtk/segmentation (tag brain bounti tissue).

## 1. Introduction

Fetal MRI provides complementary diagnostic information to antenatal ultrasound (Rutherford et al. (2008)) and allows detailed characterisation of normal and abnormal patterns of fetal brain development based on both visual analysis and quantitative metrics. Dedicated acquisition protocols for structural fetal MRI such as single shot turbo spin echo (SSTSE) and recent developments in retrospective motion correction methods such as 3D slice-to-volume registration (SVR) have lead to the generation of high-resolution 3D isotropic images Prayer et al. (2017); Aertsen et al. (2020)Gholipour et al. (2010); Kuklisova-Murgasova et al. (2012); Ebner et al. (2020)Uus et al. (2022). The continuity of these images in 3D space allows for accurate 3D segmentation and volumetry of individual brain Regions-of-Interests (ROIs) or tissue compartments. The protocols for measurements and segmentation of brain structures (relevant to a specific study) have been commonly defined in a reference space represented by either individual subject datasets at different gestational ages (GA) or population-averaged atlases Gholipour et al. (2017).

In fetal MRI studies, manual brain segmentations are conventionally performed in 2D planes on 3D T2w SVR-reconstructed images. This is a time consuming process and manual labels are generally prone to errors including inconsistencies in through plane views, missing finer parts of large region of interest (ROIs) (e.g., cortex) and over- or underestimation at tissue interfaces.

The classical automated brain parcellation methods widely used for fetal MRI rely on simple single or multiatlas registration-guided label propagation with probabilistic label fusion. However, these methods are notably prone to significant under- / over-estimations at tissue interfaces due to limits of registration (especially for the cortex). As a result, the most recent volumetric fetal brain studies Story et al. (2021); Rollins et al. (2021); Machado-Rivas et al. (2021); Vasung et al. (2022) reported the need for substantial manual refinement of segmentations produced by automated atlas-based methods.

In contrast, the results of the recent FETA fetal MRI challenge Payette et al. (2021) suggest that deep learning provides robust performance for multi-label segmentation in T2w 3D SVR images. Here, commonly used baseline models included conventional 2D and 3D UNet convolutional neural networks (CNN) Ronneberger et al. (2015); Özg ü n Çiçek et al. (2016), as well as use of the more recent advanced nnUNet Isensee et al. (2021). Khalili et al. (2019) proposed one of the first works that used 2D UNet for brain tissue parcellation in 2D fetal MRI slices. Since then, a number of works focused on different challenges specific to fetal MRI. For example, Fidon et al. (2021a) introduced label-set loss functions for cases with partial input annotations, and Fidon et al. (2021b) proposed distributionally robust optimisation to improve generalisation for unseen abnormal cases with anatomical variabilities. Other works by Li et al. (2021); Pei et al. (2021) used anatomical priors to train conditional atlases and increase performance. Deep attentive modules Dou et al. (2021) and incorporation of additional topological information de Dumast et al. (2021); Li et al. (2022) were used to improve cortical segmentation. Recently, Karimi et al. (2023) also reported a significant improvement of segmentation results based on training on smoothed noisy segmentations of globally refined propagated atlas labels Gholipour et al. (2017).

One of the remaining major challenges is caused by the lack of established high quality ground truth segmentations. Even with advanced deep learning methods, use of low quality training labels is generally expected to propagate systematic errors to the predicted segmentation and volumetry outputs. This is made more challenging, since there is still no universally accepted reference parcellation protocol for fetal brain anatomy. Recent quantitative studies relied either on publicly available atlas parcellation maps Gholipour et al. (2017) or internal fetal MRI exper-115 tise. Reliability if further affected by different acquisition parameters, image quality and anatomical variations.

## 2. Contributions

In this work, we propose a new refined protocol for parcellation of fetal brain tissue in 3D T2w SVR-reconstructed fetal brain images. It is defined a set of labels for the T2w channel of the new publicly available 21-36 GA fetal brain MRI atlas Uus et al. (2023) from the developing Human Connectome Project (dHCP). This protocol is then used as a basis for semi-supervised training of a dedicated deep learning segmentation pipeline (BOUNTI) for 3D T2w SVR-reconstructed images. The training of networks is based a large set of high-quality brain segmentations created by thorough manual refinement of labels propagated from the GA-matched atlases. In addition, the feasibility of the pipeline is assessed by comparison of brain tissue volumetry growth charts created from segmentations of 390 3D T2w brain images from three normal cohorts, acquired with different acquisition protocols, and quantitative volumetric comparison between 60 normal control and 65 ventriculomegaly cases.

## 3. Methods

This work proposes a practical deep learning segmentation solution for 3D T2w fetal brain images for automated volumetric analysis of large cohorts that would minimise the need for excessive manual editing. Fig. 1 summarises the main components of implementation of the proposed pipeline. At first, we define a new refined brain tissue parcellation protocol in the dHCP fetal atlas space. The atlas label propagation in combination with thorough manual refinement is then used to generate a large consistent set of high quality segmentations of 3D SVR fetal brain images from cohorts with different acquisition protocols. The brain segmentation networks are trained in several iterations based on semi-supervised approach.

**Figure 1:**
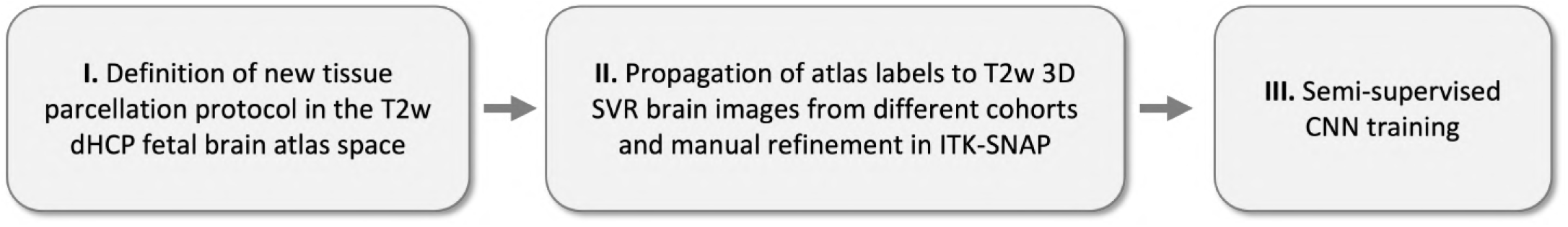
The main steps of the proposed solution for implementation of a deep learning pipeline for 3D brain tissue parcellation for motioncorrected 3D fetal MRI.

### 3.1. Cohorts, datasets and preprocessing

The fetal brain T2w SSTSE MRI datasets used in this work were acquired as part of different studies at Kings College London with different acquisition protocols. These included:

- 302 fetal participants from the developing Human Connectom Project - dHCP (REC 14/Lo/1169) project acquired on 3T Philips Achieva MRI system with a 32-channel cardiac coil using a dedicated dHCP fetal acquisition protocol Price et al. (2019) with TE=250ms, acquisition resolution 1.1 × 1.1mm, slice thickness 2.2mm, -1.1mm gap and 6 stacks;
- 55 fetal participants from the Intelligent Fetal Imaging and Diagnosis - iFIND (REC 14/LO/1806) project acquired on 1.5T Philips Ingenia MRI system using 28-channel torso coil with TE=80ms and TE=180ms, acquisition resolution 1.25×1.25mm, slice thickness 2.5, -1.25mm gap and 9-11 stacks;
- 85 fetal participants with cardiac anomalies from the fetal CMR service at Evelina London Children’s Hospital (REC 07/H0707/105) acquired on 1.5T Philips Ingenia MRI system using 28-channel torso coil with165 TE=80ms, acquisition resolution 1.25 × 1.25mm, slice thickness 2.5, -1.25mm gap and 9-11 stacks;
- 91 fetal participants from the Placental Imaging Project-PiP (REC 16/LO/1573) and Individualised Risk prediction of adverse neonatal outcome in pregnancies that deliver preterm using advanced MRI techniques and machine learning study (REC 21/SS/0082), on 3T Philips Achieva MRI system using a 32-channel cardiac coil with TE=180ms, acquisition resolution 1.25 × 1.25mm, slice thickness 2.5, -1.5mm gap and 5-6 stacks;
- 55 fetal participants from the CARP (REC 19/LO/0852) and Placental Imaging Project -PiP (REC 16/LO/1573) projects acquired on 1.5T Philips Ingenia MRI system using 28-channel torso coil with TE=180ms, ac-SVR method to 0.75-0.8mm isotropic resolution using the quisition resolution 1.25 × 1.25mm, slice thickness 2.5, -1.25mm gap, and 4-5 stacks;
- 125 fetal participants from the Quantification of fetal growth and development using magnetic resonance imaging study (REC 07/H0707/105) acquired on 1.5T Philips Achieva MRI system using 32-channel cardiac coil with TE= 160ms, acquisition resolution 1.25 × 1.25mm, slice thickness 2.5, -1.5mm gap and 8 stacks.

The cohort includes both normal control and a subset of cases with abnormal findings but without extreme deviations in their anatomy (presence and preserved global shape of all brain regions and absence of lesions). The GA of participants varies within 20-38 weeks range.

All 3D brain images were reconstructed using fully automated SVR motion-corrected pipelines. The dHCP datasets were reconstructed using the dedicated dHCP SVR method Cordero-Grande1 et al. (2019) to 0.5mm resolution and reoriented Wright et al. (2018) to the standard radiological space. The rest of the datasets were reconstructed based on the Kuklisova-Murgasova et al. (2012) automated version^2^ that also includes automatic reorientation to the standard space. We used only acceptable quality datasets with sufficient visibility of all brain structures and anatomical features. SVR reconstructions with intensity artifacts or failed motion correction (that on average tend to be present in approximately 10-15% of all cases) were not included in this study.

### 3.2. Proposed brain tissue parcellation protocol

Currently, there is no established consensus in the existing fetal brain manual annotation protocols (e.g., Gholipour et al. (2017); Kyriakopoulou et al. (2017); Khalili et al. (2019); Payette et al. (2021)). This is further complicated by the difference between the fetal and neonatal anatomy and lower image resolution of fetal MRI (e.g., DRAW-EM Makropoulos et al. (2018)) relative to the size of the fetal brain. Therefore, we formalised a new brain tissue segmentation protocol that is defined as a set of labels in the new fetal brain MRI dHCP atlas (Uus et al. (2023)) space using the T2w channel and 16 timepoints from 21 to 36 weeks.

The inclusion and definition criteria for individual tissue structures were based on the clinical relevance to quantitative studies, evidence from the anatomy histology atlases Bayer and Altman (2003, 2005) and the clear visibility of the structures in individual subject 3D T2w SVR MRI images. At first, we used the optimised neonatal dHCP DRAW-EM pipeline Makropoulos et al. (2018) to create an initial set of segmentations of the major tissue structures. Next, a clinical researcher (VK), with more than 10 years experience in fetal MRI, manually refined and subdivided (when relevant) the labels to 19 ROIs using ITK-SNAP^3^ including cortical grey matter (GM), white matter (WM), cerebrospinal fluid (CSF), deep GM (DGM), ventricles, cavum, brainstem and cerebellum ROIs. The refinement was based on the fetal brain anatomy guidebooks Bayer and Altman (2003, 2005) and performed for 3 atlas timepoints: 25, 30 and 36 weeks GA. This was followed by left/right separation of all paired structures. These preliminary parcellation maps were propagated to the rest of the atlas timepoints using nonlinear registration from MIRTK^4^ and manually refined, when required. These parcellations were then used to generate the ground truth labels for training of the proposed BOUNTI segmentation network described in the next section. The final set of atlas parcellation maps was created by using the final version of the trained BOUNTI segmentation pipeline (described in Sec. 3.3). This was done to reduce any partial volume effects at the cortex interface or inconsistencies in manual segmentations.

The atlas along with the proposed tissue parcellation maps is publicly available at the online Centre for the Developing Brain (CDB) data repository ^5^.

### 3.3. Automated parcellation of the fetal brain BOUNTI segmentation pipeline

The proposed pipeline for “Brain vOlumetry and aU-tomated parcellatioN for 3D feTal MRI” (BOUNTI) is summarised in Fig. 2.A. It consists of two main steps.

**Figure 2:**
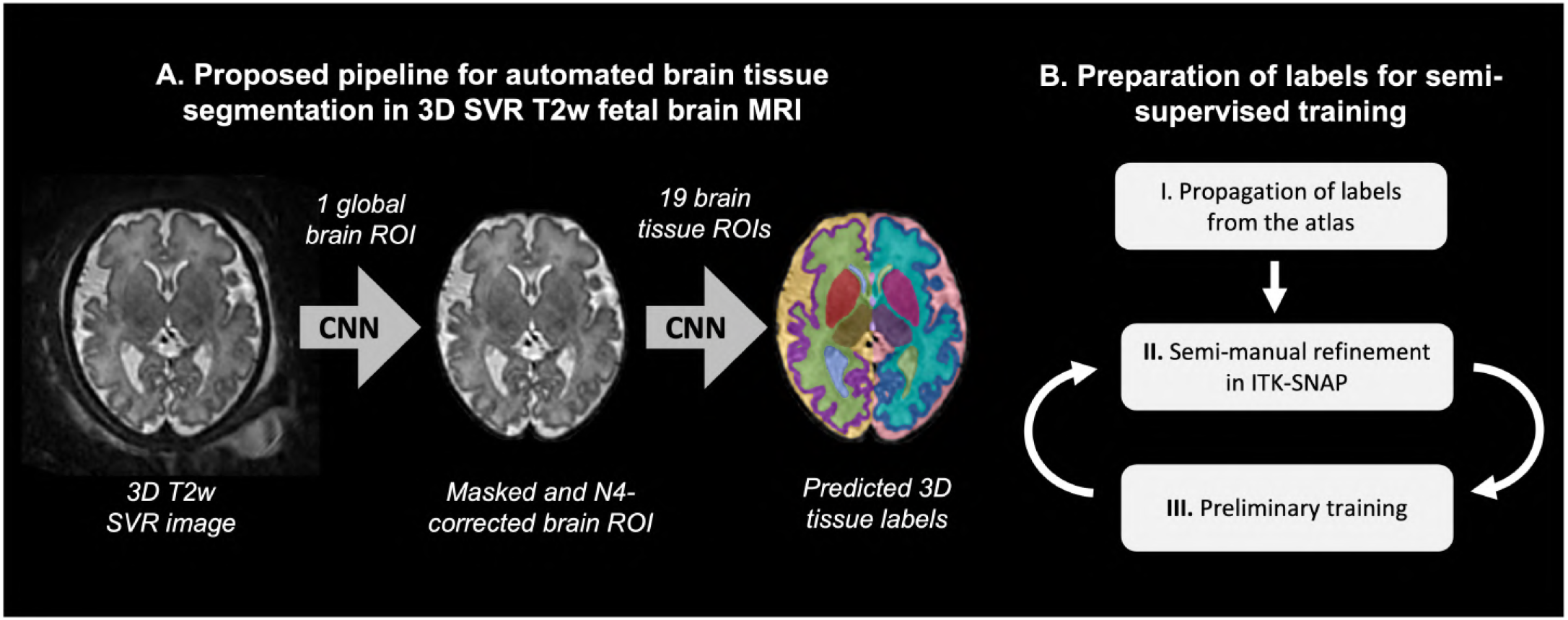
Proposed BOUNTI pipeline for automated brain tissue parcellation for 3D T2w fetal MRI (A) and preparation of labels for semi-supervised training (B).

Initially, a 3D CNN module is used for global localisation of the brain in order to remove any present external background ROI. This also increases robustness to variations in formats of outputs of different SVR methods (e.g., with or without masking and padding). Next, the cropped, masked and bias corrected 3D brain ROI is segmented using a multi-label 3D CNN module trained on datasets with manually refined labels propagated from the atlas using registration. Taking into account the large number of datasets required for training, as well as the time-consuming nature of the manual refinement process, we employed a semi-supervised approach with several iterations. At each iteration, a preliminary version of a pretrained network is used for generation of a larger dataset followed by manual fine-editing of labels for the final training stage.

#### Selected deep learning models

For the segmentation pipeline, we chose to use a combination of a classical 3D UNet Özgün Çiçek et al. (2016) and an attention UNet Oktay et al. (2018) architectures for both CNN modules for brain extraction and tissue parcellation. While 3D UNet is a robust and well established tool widely used in 3D fetal MRI Payette et al. (2021), attention UNet reportedly improves prediction performance by focusing on important structures of varying shapes.

We used the standard MONAI Cardoso et al. (2022) 3D UNet and Attention-UNet implementations with five and four encoder-decoder blocks (output channels 32, 64, 128, 256 and 512), correspondingly, convolution and upsampling kernel size of 3, ReLU activation, dropout ratio of 0.5. We employed AdamW optimiser with a linearly decaying learning rate, initialised at 1×10^*−*3^, default *β* parameters and weight decay=1×10^*−*5^. The input image dimensions are 128×128×128 and 256×256×256 and the outputs have 2 and 20 channels (with background) for global localisation and tissue parcellation networks, correspondingly. At inference, the predictions (per tissue ROI) of 3D UNet and Attention-UNet networks are averaged at the output before softmax. In addition, in order to improve robustness of the pipeline and limit any possible bias in segmentation of the left and right structures, the images are passed through the network twice, in the original orientation and flipped along Y-axis. The second prediction is then flipped to the original orientation, and order of the output channels is updated. The two segmentations are then averaged to produce the final output.

#### Selected datasets for training and testing

The training for both brain extraction and tissue parcellation was performed on datasets with different acquisition protocols and SVR reconstruction methods. The summary is provided in Tab. 1. The GA range of the participants varies within the 21-38 weeks GA range. The testing dataset includes 40 randomly selected (not used in training) 3D T2w SVR reconstructed images from 4 different acquisition protocols.

**Table 1:** Summary information about training and testing datasets.

| Cohort | Acquisition | 3D SVR reconstruction |
| --- | --- | --- |
| *Training datasets* |  |  |
| 190 (normal and abnormal) | 3T, TE=250ms | Cordero-Grande et al. (2019)) |
| 120 (normal and abnormal) | 1.5T, TE=80ms | Kuklisova-Murgasova et al. (2012)) |
| 70 (normal and ventriculomegaly) | 1.5T, TE=160/180ms | Kuklisova-Murgasova et al. (2012)) |
| *Testing datasets* |  |  |
| 10 (normal and abnormal) | 3T, TE=250ms | Cordero-Grande et al. (2019)) |
| 10 (normal and abnormal) | 1.5T, TE=80ms | Kuklisova-Murgasova et al. (2012)) |
| 10 (normal and abnormal) | 1.5T, TE=180ms | Kuklisova-Murgasova et al. (2012)) |
| 10 (normal and abnormal) | 3T, TE=180ms | Kuklisova-Murgasova et al. (2012)) |

#### Image preprocessing

Taking into account the varying size, resolution and intensity ranges of input SVR reconstructions, the general preprocessing steps for all images in both steps of the pipeline included: transformation to the standard radiological space coordinate system, cropping of the background, resampling with padding to the required input grid size and rescaling to 0-1 range. For the brain tissue segmentation step, the brain images were also masked and cropped with the dilated (2 iterations) global brain masks obtained from the first step of the BOUNTI pipeline. All preprocessing steps were implemented based on MIRTK toolbox.

#### Preparation of training datasets

For both CNN steps of the pipeline, training was performed in three stages. In this case, a version of the network pretrained on a small number of cases (from the previous iteration) was used for segmentation of the main training dataset. This strategy was selected for both practical reasons and required accuracy and consistency of the ground truth labels.

For the brain extraction CNN step, we used an already existing network (3D UNet in MONAI pretrained on datasets from other research projects at our institution) to segment all SVR brain reconstructions with different background coverage and masking. The output segmentations were inspected and manually refined (at every stage), when required, resulting in 380 images for final training. We augmented the training dataset by allowing image masking to be performed with a varying degree of dilation (2 to 18 iterations) of the brain mask and background padding. For the first stage of training of the brain tissue parcellation CNN, we generated labels via registration-guided atlas propagation n (19 ROIs) for a preliminary set of 200 fetal brain from dHCP and iFIND/fCMR cohorts covering 20-38 weeks GA range. Image registration was run with multi-channel (T2w and cortex generated by an existing in-house 3D UNet trained on Draw-EM cortical segmentations for dHCP cohort) nonlinear MIRTK registration Rueckert et al. (1999) with LNCC metric and 6mm window size. After visual inspection, 100 images (distributed across the whole GA range) with the minimal/average amount of errors (based on visual assessment of cortical interface) were selected and semi-manually refined by researchers (AU, DC, AD) trained in fetal MRI (Fig. 2.B). The refinement included editing of the cortex, WM, CSF, deep GM and cerebellum ROIs using the user-guided active contour segmentation tool Yushkevich et al. (2006) in ITK-SNAP, followed by local manual editing of individual tissue labels. In general, label propagation worked well for 21-31 weeks GA range with only minimal editing required (mainly for low image quality or cases with abnormal findings). For late GA (*>*34 weeks) cases, refinement required approximately 1-2 hours per case, primarily focusing on the cortex ROI. Notably, even after refinement the cortex labels were not smooth and had minor discontinuities / overlap between WM/GM/CSF ROIs. In addition, the training dataset was extended by flipping of the brain in a left/right direction (with corresponding changes of the label numbers) and histogram matching to different TE references (e.g., from TE=250ms to TE=80ms) as augmentation for the flipped versions.

This first version of the refined datasets was used for preliminary training of a 3D UNet. The network was then used to segment all 380 fetal brain images from a mixture of the available cohorts (including the originally refined) with more early (*<*22 weeks) and late (*>*34) participants to balance the extreme anatomy differences. All labels were reviewed in terms of errors and 200 images were selected for further training. The output CNN labels for the selected images were again edited in ITK-SNAP (using active contours and manual refinement), when required. The new training dataset was then used to train the network with the same data augmentation strategy followed by segmenting, review and editing of the next part of 3D brain images. This procedure was repeated for a second time. The final training dataset consisted of 380 images (doubled by flipping augmentation).

#### Training of the networks

Training of all networks was performed in MONAI framework using soft Dice and cross entropy loss and AdamW optimiser. The final training stage of the brain extraction 3D UNet and Attention-UNet was performed on 360 training and 20 validation datasets for 20000 and 50000 iterations, correspondingly. The networks were trained separately. The final training of the brain tissue parcellation 3D UNet and Attention-UNet was performed on 360 training and 20 validation datasets for 150000 and 300000 iterations, correspondingly. We used standard MONAI augmentation including: bias field, affine rotations, Gaussian noise and blurring.

#### Docker for the brain segmentation pipeline

The proposed fetal brain segmentation pipeline with trained networks is publicly available as a standalone docker application^6^ at SVRTK fetal segmentation repository. In order to ensure high segmentation quality, the main input requirements to 3D fetal MRI SVR reconstructed images include: T2w contrast (1.5/3T, TE=80-250ms), orientation in the standard radiological space, 21-38 weeks GA range, no extreme structural anomalies, sufficient image quality (in terms of clear visibility and definition of the brain structures) and no extreme SNR loss or shading artifacts.

### 3.4. Growth charts of the fetal brain development

In order to assess the practical application of the proposed dHCP brain parcellation protocol and segmentation pipeline, we used the BOUNTI pipeline for segmentation of 244 normal control fetuses from the dHCP project (185 of these datasets were used during the training stage) and 146 normal participants from two other cohorts (1.5T, TE=180ms and 3T, TE=180) with different MRI acquisition parameters from 21 to 38 weeks GA range and singleton pregnancies (these datasets were not used in training). All images were resampled to 0.5mm isotropic resolution. All segmentations were reviewed and manually fine-edited, if required, in order to ensure reliability of nomograms and assess the impact on global trends.

The label volumetry was used to create growth charts (mean, 5^th^ and 95^th^ centiles) for brain development of 9 structures (combined right/left and associated ROIs) based on the guidelines from Royston and Wright (1998). The output table-format centile calculator for all individual structures is available at the atlas repository. The statistical difference between the cohorts was assessed for the main solid tissue structures (WM, cortical GM, deep GM) using ANCOVA analysis (volume measurements were converted to log format, when relevant).

#### Comparison of normal control and VM cohorts

In addition, to test the performance of the BOUNTI pipeline we utilised imaging data from our previously published study Kyriakopoulou et al. (2017) and compared the volumetric results. We run BOUNTI on a cohort of fetuses with ventriculomegaly (65) and a normal control (60) cohort. All 125 fetal MRI datasets were reconstructed using the classical SVR method Kuklisova-Murgasova et al. (2012), reoriented to the standard space and resampled to 0.5 mm resolution. The 3D brain images were segmented using the BOUNTI pipeline and visually inspected and manually refined, when required. The volume of the total lateral ventricles and suprantentorial brain tissue was compared between the cohorts using ANOVA with and without manual refinement.

## 4. Results and experiments

### 4.1. Proposed tissue parcellation protocol

The proposed multi-tissue parcellation protocol defined for the dHCP fetal brain MRI atlas^7^ is shown in Fig. 3. It includes 19 major brain tissue ROI labels: cortical GM, fetal WM, external CS, lateral ventricles, cavum, thalamus, basal ganglia, brainstem, cerebellum, vermis, 3rd and 4th ventricles, with left/right separation of paired structures. Fig. 4 demonstrates parcellation maps at 21, 26, 31 and 36 weeks that reflect the expected changes during normal brain development such as cortical folding and shape of the ventricles. Rendered cortical GM and WM have smooth boundaries without gaps and with well defined cortical folds. The cortical ribbon parcellation is thin without partial volume effect in WM.

**Figure 3:**
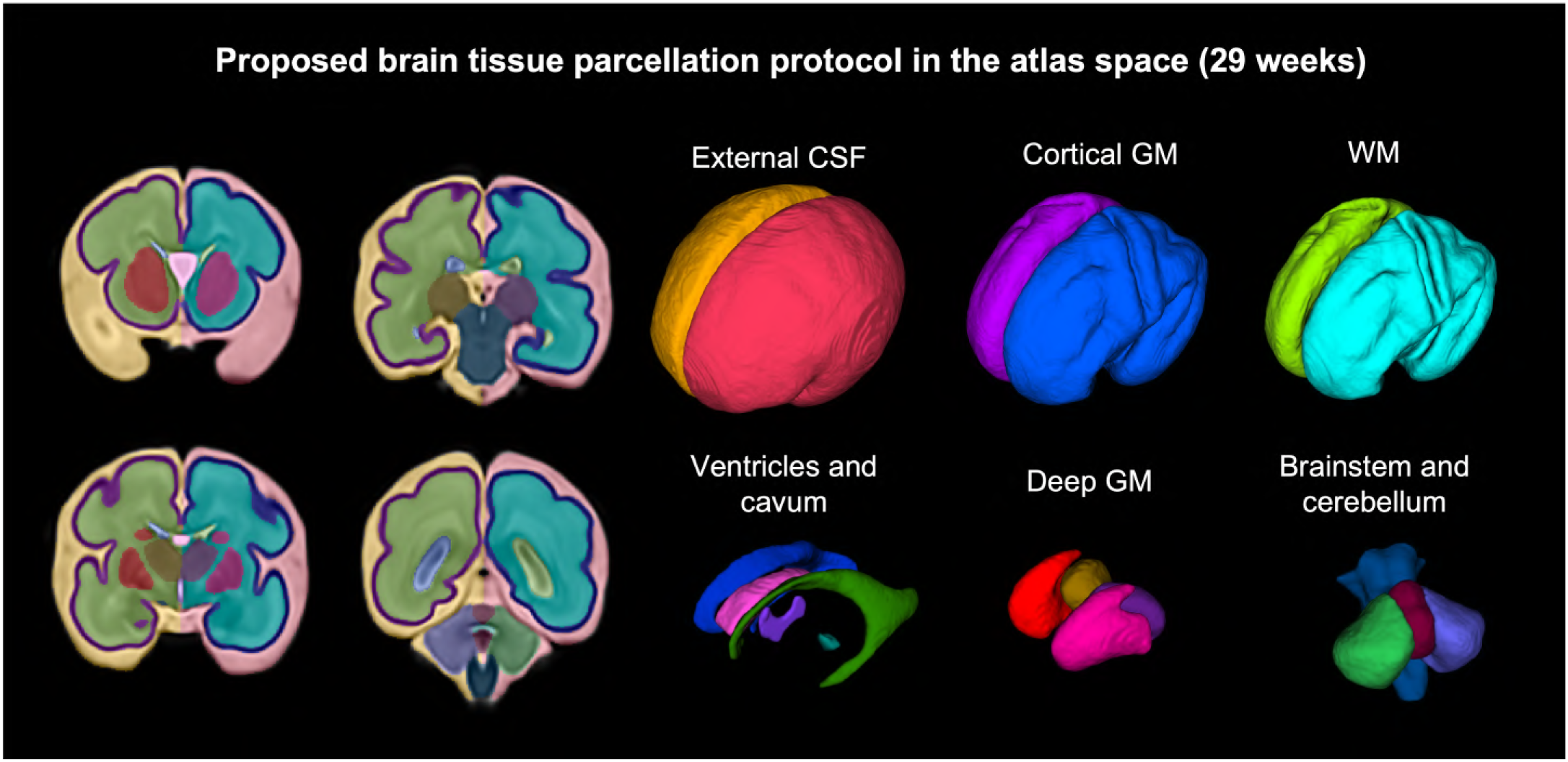
Proposed tissue parcellation protocol defined in the dHCP fetal brain MRI atlas.

**Figure 4:**
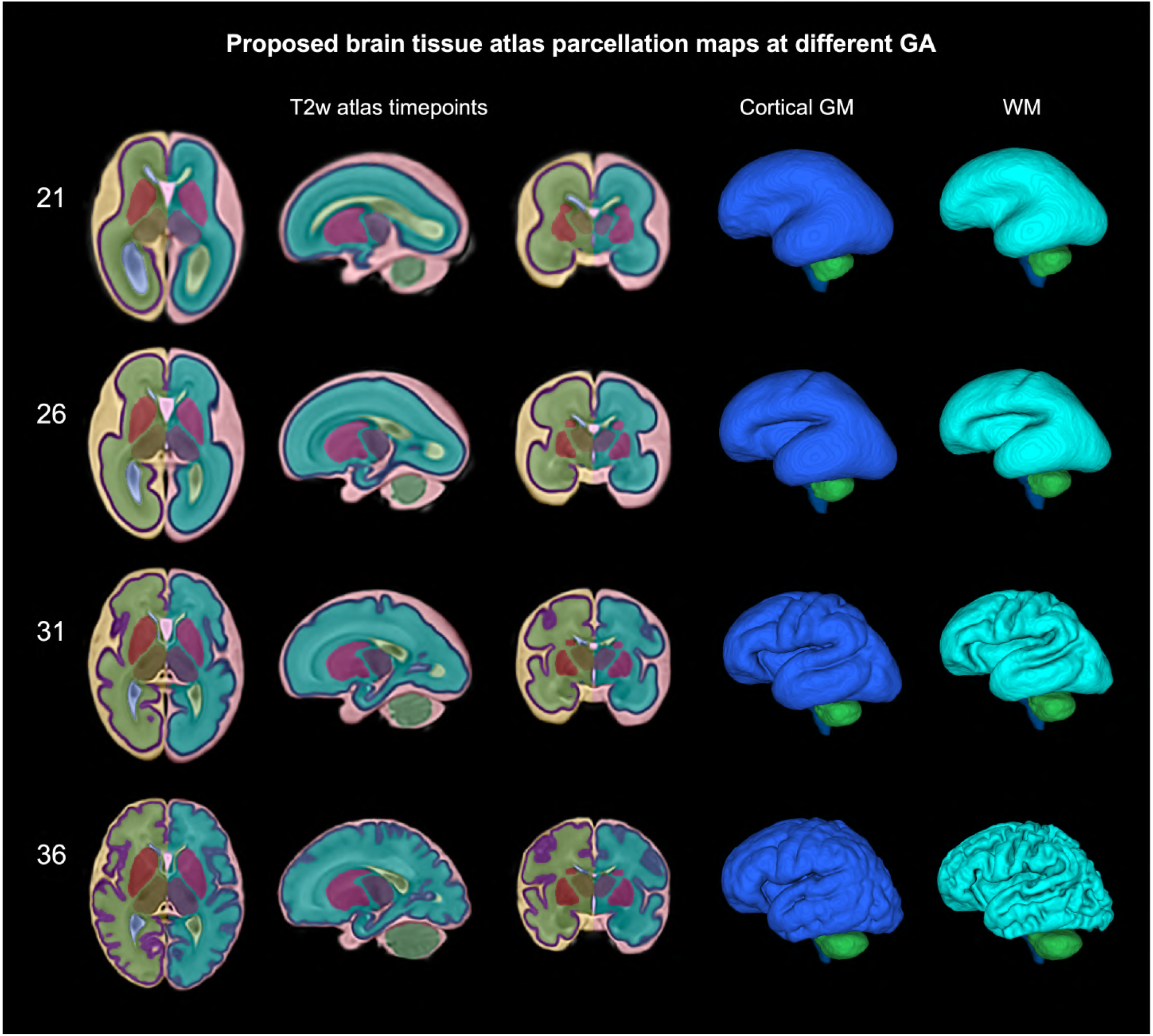
Proposed multi-tissue brain parcellation maps at 21, 26, 31 and 36 weeks GA timepoints of the dHCP fetal brain MRI atlas.

### 4.2. Automated parcellation of the fetal brain Comparison with semi-manual GT segmentations

The results of testing of the BOUNTI pipeline on 40 T2w SVR brain images from 4 different acquisition protocols and 21-38 weeks GA range (Tab. 2) showed robust performance for all tissue ROIs. In all 40 test cases, all structures were correctly globally detected in all BOUNTI segmentations (100 % detection rate). The relatively high Dice values are expected due to the high quality and consistency of training datasets generated by label propagation and thorough manual refinement. There were no systematic differences in the method performance for different test groups apart from slightly higher Dice values for the dHCP cohort (3T, TE=250ms). This potentially is due to higher tissue contrast and spatial resolution of the images, as well as similarity to the atlas. The relative volume differences are within the generally acceptable range. Even after manual and active contour refinement, the classical registration-based LP is prone to minor inconsistencies for the cortex ROI especially at late GA due to complex folding patterns. Fig. 5 demonstrates that BOUNTI corrected overestimation of CSF and cortical GM that were present in the semi-manual ground LP labels. The CNN outputs have significantly smoother cortex boundaries and less errors at different tissue interfaces than the ground truth LP. This is in agreement with the previously reported successful CNN performance in the recent FETA brain MRI challenge Payette et al. (2021).

**Figure 5:**
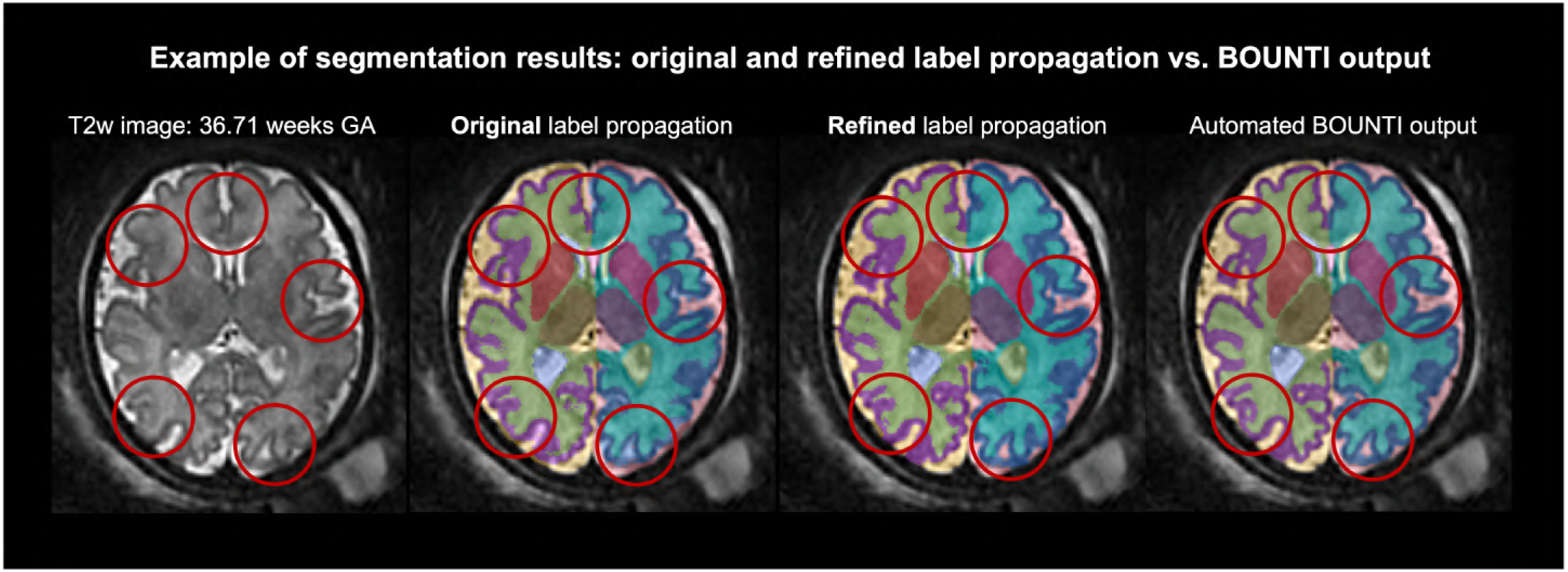
An example of comparison of BOUNTI segmentation results vs. the original and manually refined (ground truth) label propagation: late GA test case from dHCP cohort (3T, TE=250ms).

#### Impact of GA at scan

Fig. 6 shows BOUNTI segmentation results for test cases at different GA ranges. In general, the network performed better for the early and medium GA ranges. Several minor errors were present at the back of the cortex ROI in 6 late GA cases *≥*35 weeks. This suggests the further need for incorporation of topological information Li et al. (2022) to ensure spatial continuity of individual structures, which will be especially relevant for surface-based analysis as well as late GA cases.

**Figure 6:**
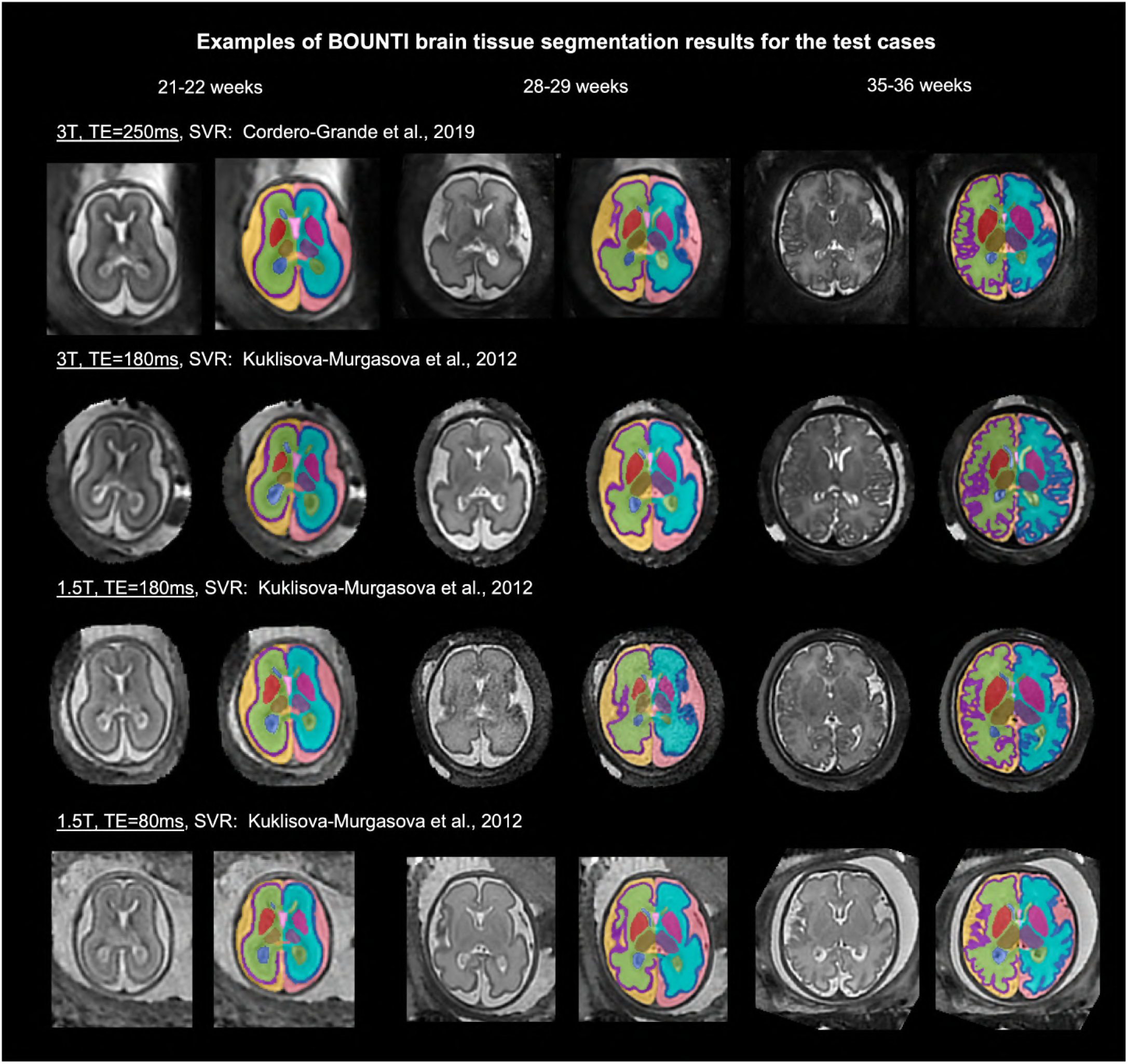
Examples of the outputs of segmentation pipeline for the test cases with different acquisition protocols and gestational ages.

#### Impact of acquisition and SVR reconstruction parameters

The examples of the segmentation results for all four acquisition protocols with 1.5T and 3T field strength and 80, 180 and 250ms echo time in Fig. 6 show clearly different tissue contrasts and different image quality in terms of definition of the finer features (due to SNR levels and blurring). In addition to the quantitative analysis (Tab. 2), visual assessment of all test cases did not reveal distinct differences in terms of the network performance. This is also in agreement with the experiments in Payette et al. (2020) that the choice of SVR methods does not significantly affect segmentation performance (excluding cases with failed reconstruction).

#### Impact of image quality and artifacts

Similarly to acquisition parameters, intensity artifacts or low SNR alter tissue contrast and visibility of structures, which in turn tend to affect accuracy and certainty of segmentations. The examples in Fig. 7.A show the results of BOUNTI segmentation pipeline for previously unseen suboptimal image quality cases. While the network seemed to produce relatively stable results for low SNR regions, it failed in the cortex ROI with severe B1 shading (that could not be resolved by N4). This suggests that the degree of MONAI bias field augmentation used during the training was not sufficient, and more severe simulated shading artifacts (or more advanced bias field correction approach) would be required to address this issue. However, volumetry derived from low image quality datasets cannot be considered reliable by definition. Therefore, clinical translation of the pipeline would require a detailed specification of the image quality requirements and an additional step for automated assessment of expected segmentation certainty.

**Figure 7:**
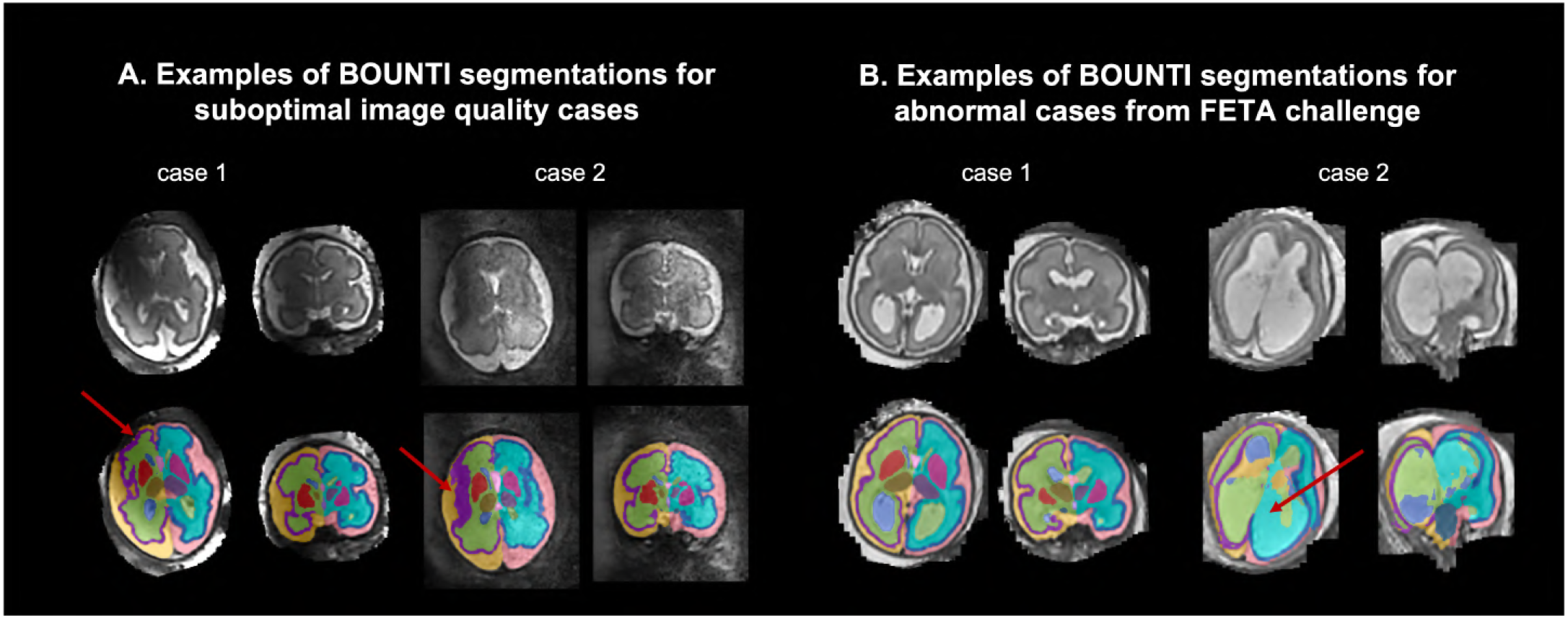
Examples of BOUNTI segmentation results for suboptimal image quality (A) and abnormal Payette et al. (2021) (B) cases.

#### Impact of anomalies

Two examples of BOUNTI segmentations for the previously unseen abnormal cases from the FETA challenge Payette et al. (2021) are shown in Fig. 7.B. Since the training dataset included abnormal cases with ventriculomegaly, visual analysis confirms that the network performance was relatively acceptable for cases with slightly enlarged ventricles. However, as expected, the network produces various errors for anomalies with significant alternations in the brain anatomy such as extreme ventriculomegaly. The optimal solution would require further training on large abnormal cohorts (e.g., similarly to Fidon et al. (2021b)) along with possible introduction of classification step and optimisation of the network architecture and parcellation protocol. Notably, the best reported performances in terms of average Dice from the FETA segmentation challenge Payette et al. (2022) vary within 0.78-0.79 range (vs. 0.87-0.90 for BOUNTI performance540 Tab. 2), which highlights the advantages of using high quality consistent ground truth labels for training.

#### 4.2.1. Alternative fetal brain segmentation methods

Taking into account the intrinsic differences in parcel-545 lation protocols (e.g., different exclusion / inclusion strategies for cortical and deep grey matter ROIs), the outputs of BOUNTI cannot be directly quantitatively compared to alternative segmentation methods. The example of visual comparison with a classical 3D UNet trained on the original FETA Payette et al. (2021) datasets (7 ROIs) and label propagation from a GA-matched alternative atlas Gholipour et al. (2017) (27 ROIs) is shown in Fig. 8. These two datasets at 24 and 35 weeks GA are from 3T, TE=180ms cohort. In general, BOUNTI provides more robust performance for the cortex ROI with thinner and smoother cortical ribbon and smaller amount of errors in comparison the other methods. This is caused by failed registration or inconsistencies in training datasets.

**Figure 8:**
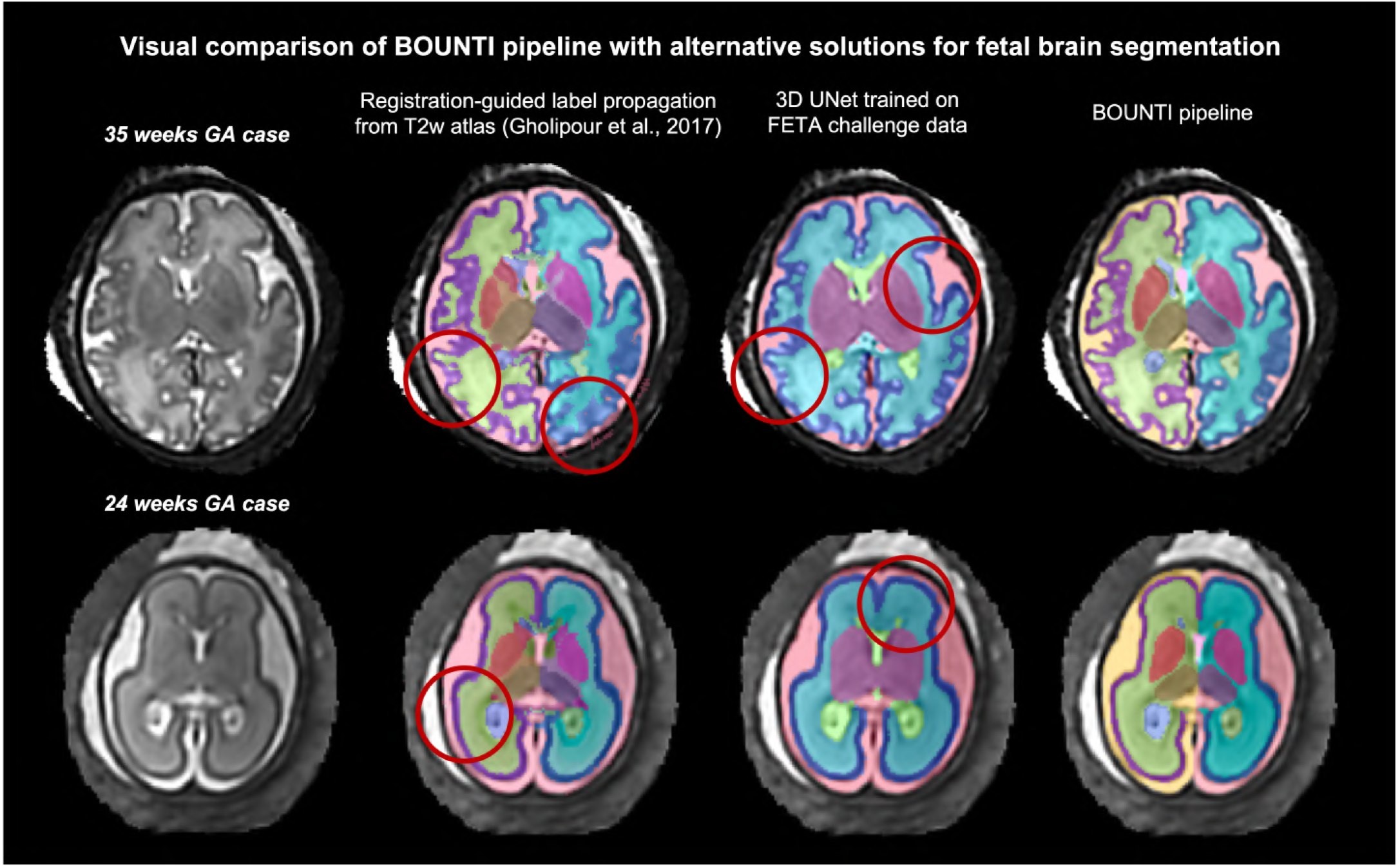
Examples of visual comparison of BOUNTI results with classical 3D UNet trained on the original FETA Payette et al. (2021) datasets with manual labels (7 ROIs) and label propagation from an alternative atlasGholipour et al. (2017) (27 ROIs): two datasets at 24 and 35 weeks GA are from 3T, TE=180ms cohort. Note: the label colours were adapted for optimal comparison.

### 4.3. Growth charts of normal fetal brain development

In order to assess the general applicability of the proposed brain atlas parcellation protocol and BOUNTI pipeline, we used it to create volumetry growth charts (Fig. 9) of the typical fetal brain development during 21 to 38 weeks GA range based on healthy control datasets from four studies with different acquisition protocols. It includes: 55 cases with 1.5T, TE=180ms (PiP, CARP); 91 cases 3T, TE=180ms (PiP, PRESTO) and 244 cases with 3T, TE=250ms (dHCP). Only cases without reported anomalies and with good quality images were selected. In order to ensure the validity of nomograms all segmentations generated by the BOUNTI pipeline were visually inspected.

**Table 2:**
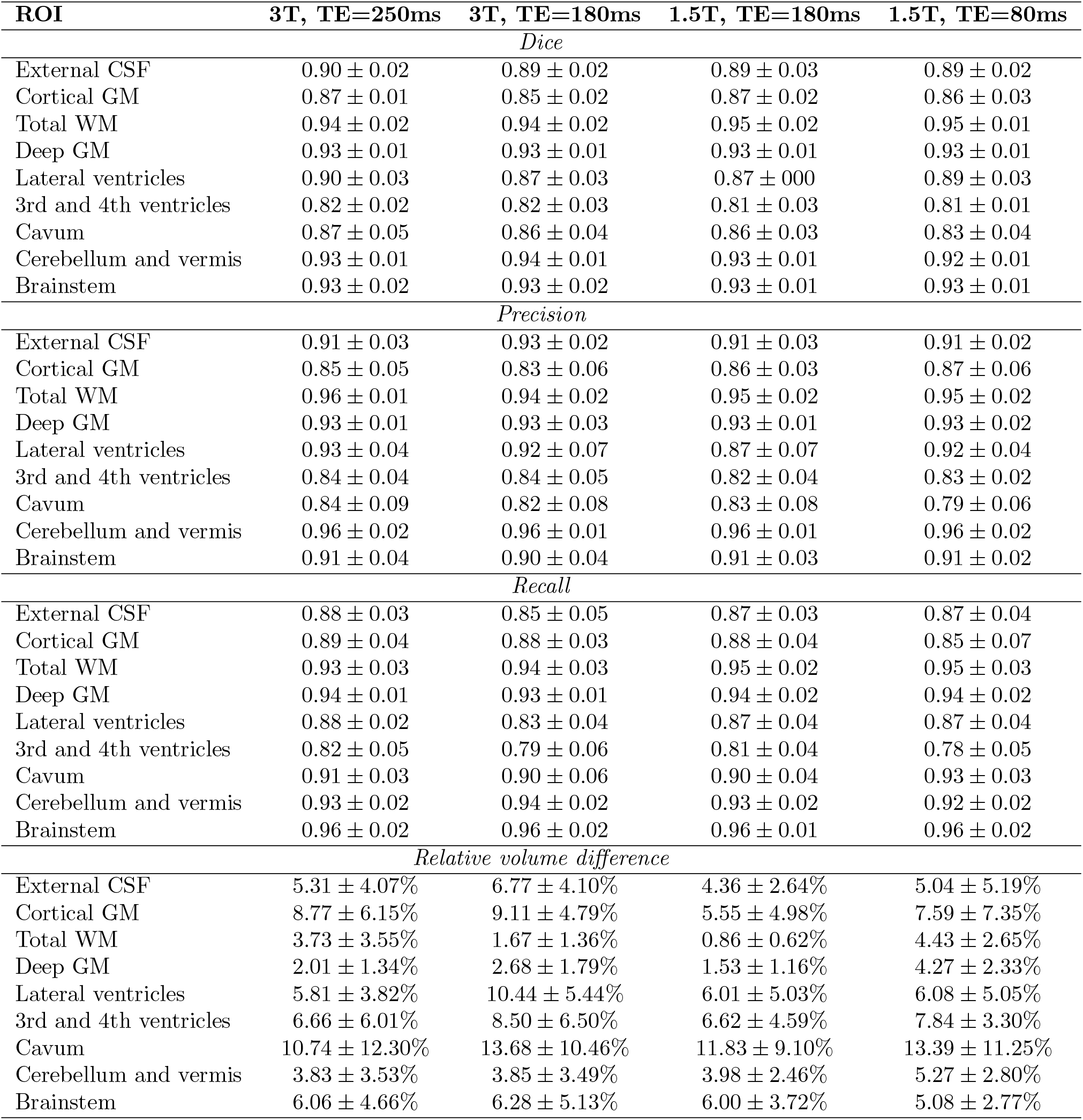
Quantitative comparison of BOUNTI segmentation results with semi-manual (label propagation with manual refinement) ground truth labels: average Dice, recall and precision. The datasets include 40 cases from 21-38 weeks GA range from 4 different acquisition protocols. Note: the labels for pair/related structures were combined.

**Figure 9:**
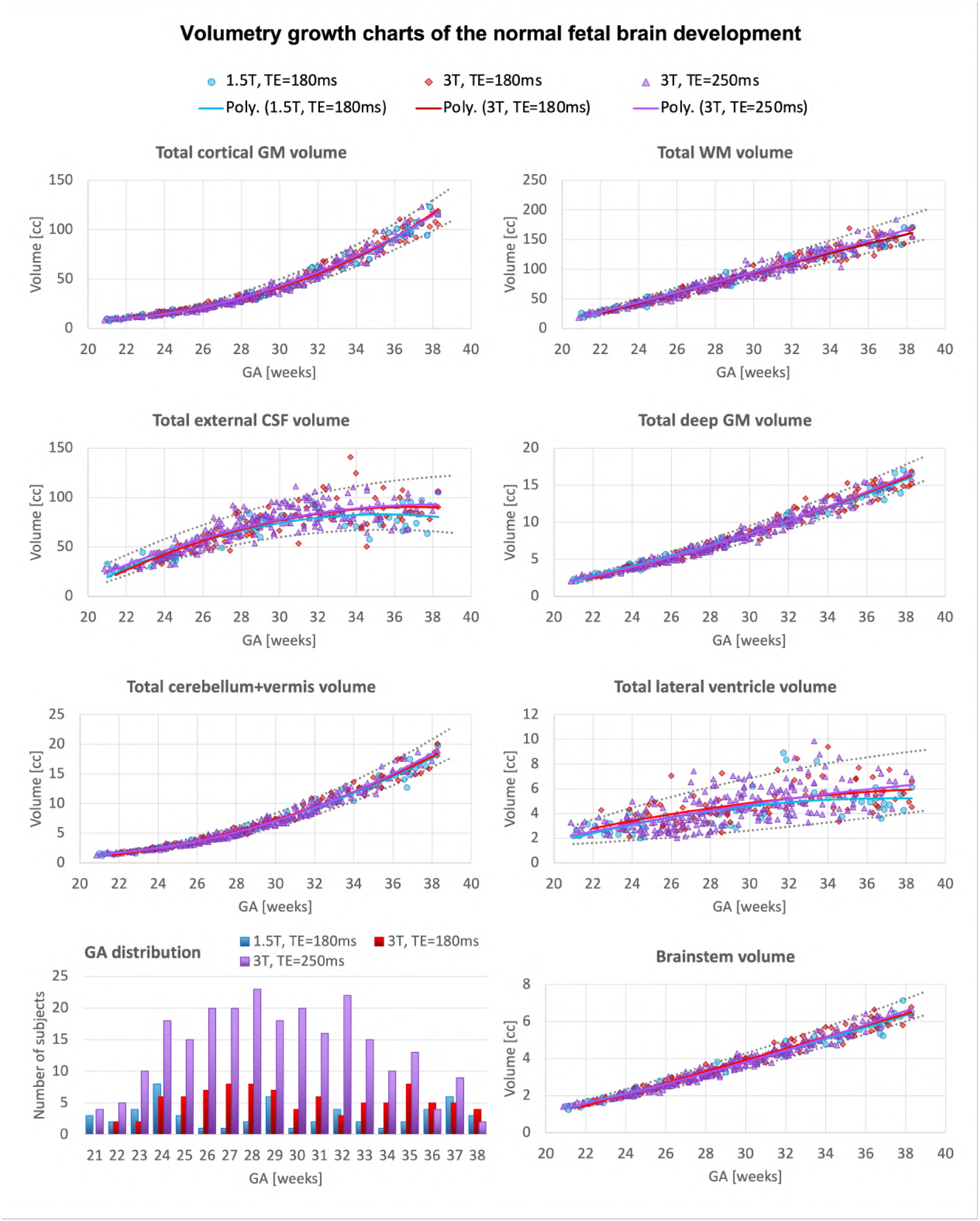
Growth charts: volumetry of the major brain structures for normal fetal cohorts from fetal MRI research studies with different acquisition parameters: 1.5T, TE=180ms (blue, 55 participants); 3T, TE=180ms (red, 91 participants); 3T, TE=250ms (lilac, 244 participants). Note: the left and right labels for pair structures were combined.

Only minor refinements at the cortex interface were required for 6 early (*<*23 weeks) and 38 late (*>*35 weeks) GA cases due to either suboptimal regional image quality or complex cortical patterns. Notably, the refinements did not produce significant changes in volumetry results (not exceeding 5% for individual labels volumes) and, on average, required less than 1-5 minutes per case of manual editing. This is a significant improvement in comparison to the previously reported required time-consuming extensive manual refinement (e.g., 1-3 hours per case in Story et al. (2021)). This further emphasises that results of any automated segmentation methods should always be inspected and confirmed before any quantitative analysis. Further development of the BOUNTI pipeline would also require integration of automated image and segmentation quality control steps as well definition of what constitutes a correct segmentation and the acceptable levels of accuracy.

The trends for all structures demonstrate the expected increase in volume with GA Kyriakopoulou et al. (2017); Machado-Rivas et al. (2021) with higher variability for external CSF and lateral ventricle volumes. There are no visible systematic deviations in the trends for different field strength and echo time. This was further confirmed by ANCOVA analysis that showed no significant differences in volumetry for the major solid tissue structures. These preliminary results show general feasibility of automated CNN solution for the accurate brain volumetry, even in the presence of acquisition differences. However, any quantitative volumetry analysis of datasets from different studies would still require careful analysis of systematic differences and implementation of a dedicated harmonisation solution.

## 4.4. Comparison on normal control and VM cohorts

Following visual inspection of the 125 datasets for any miss-labelled voxels, 6 out of the 125 cases required manual refinement in the lateral ventricle and cortical ROIs. The graphs in Fig. 10 demonstrate the volumetric comparison between the ventriculomegaly (65) and normal control (60) fetal MRI datasets (without manual editing). For this study, manual editing in the ventricle and cortex ROIs was required for 6 cases. Prior to manual editing, there is a significant difference between the cohorts, with higher supratentorial brain (*p <* 0.0001) and lateral ventricle (*p <* 0.0001) volumes in the VM cohort. The statistical significant difference in both the lateral ventricular volume (*p <* 0.0001) and supratentorial brain (*p <* 0.0001) remained following manual editing of the 6 cases. This is also in agreement with the originally reported results that were based on exclusively manually performed segmentations Kyriakopoulou et al. (2017).

**Figure 10:**
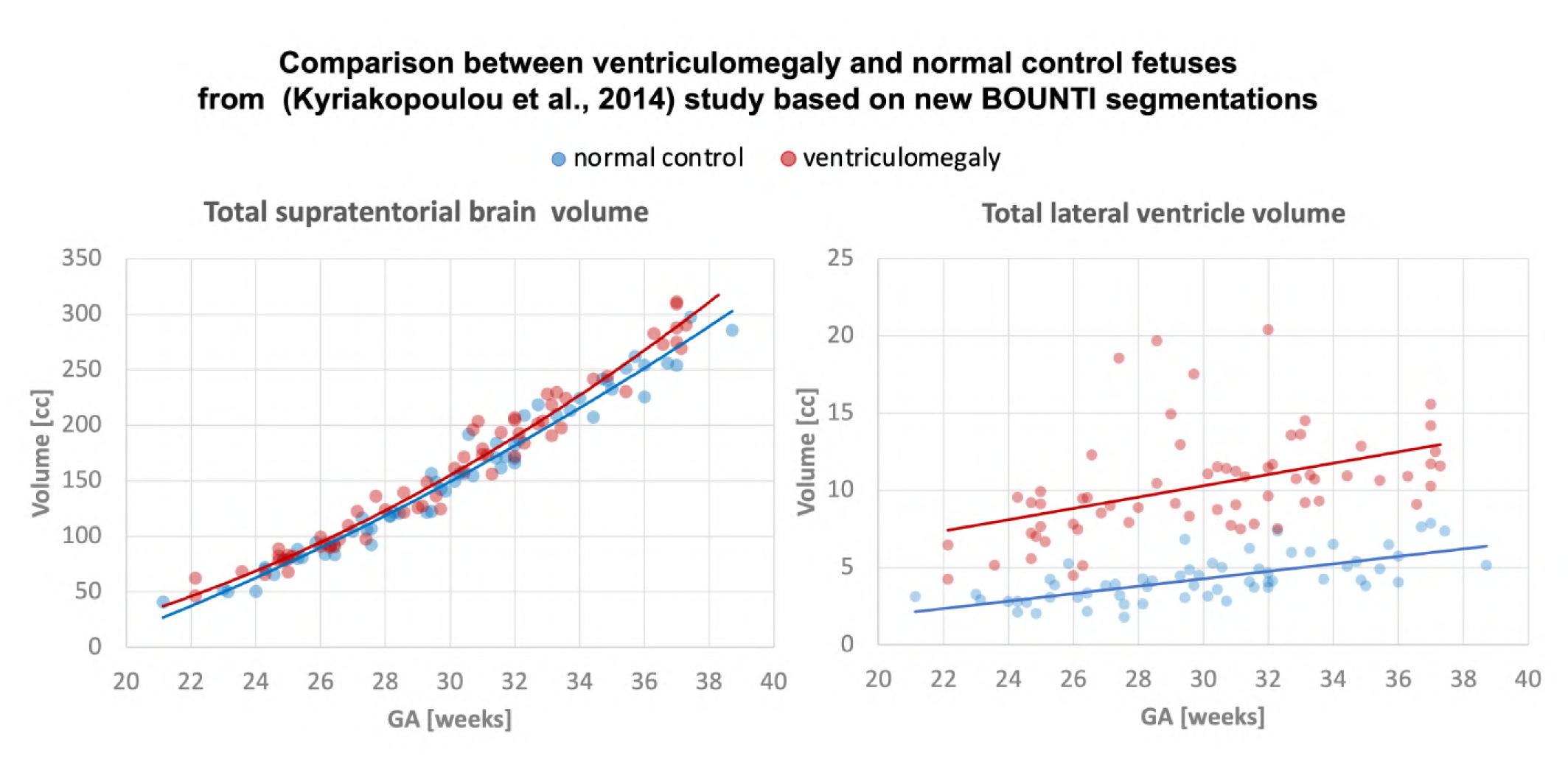
Comparison of the VM (65) and normal control (60) fetal MRI cases from Kyriakopoulou et al. (2014) study based on the new set of unedited parcellations from BOUNTI: lateral ventricle and supratentorial brain volumes. Note: the left and right labels for pair structures were combined.

## 5. Discussion

The main aim of the new dHCP tissue T2w parcellation protocol and training of the “BOUNTI” pipeline was implementation of a foundation automated multi-label segmentation tool suitable for robust and practical volumetric analysis at wide GA ranges and different acquisition protocols that would minimise required manual refinement.

First, we defined a new protocol for brain tissue parcellation in the fetal MRI dHCP atlas space for 21-36 weeks GA timepoints. The proposed parcellation map includes 19 tissue ROIs defined in the atlas space.

The CNN pipeline (based on combination of 3D Attention and classical UNet) was trained on 360 T2w 3D SVR-reconstructed brain images from 6 studies with different acquisition protocols and SVR reconstruction methods. In order to generate a high quality training dataset, we used a semi-supervised approach based on several iterations of manual refinement of atlas label propagations and outputs of the pretrained network. The results of testing on 40 images from different cohorts showed consistent performance for the whole GA range and varying image contrast. The predicted segmentations have a well defined cortex interface with only minor errors present in a small proportion of late GA cases. This significantly minimises required manual editing in comparison to the alternative solutions. Furthermore, BOUNTI segmentation is significantly faster (1-5 minutes per case depending on system configuration and image parameters) than the alternative classical methods such as Draw-EM or label propagation.

The BOUNTI pipeline was then used to segment 390 3D T2w fetal brain images of the normal participants from the dCHP and two other cohorts acquired at different field strength and with different echo time. Notably, only minor manual editing was required in *<* 15% of all cases without significant changes in volumetric measurements. The corresponding generated growth charts showed no significant differences in trends of the main solid brain tissue ROIs for different acquisition parameters. Furthermore, comparison of BOUNTI label volumes between the normal and cases with VM from our earlier study Kyriakopoulou et al. (2014) showed similar significant differences. These preliminary results potentially suggest the suitability of using one universal network for segmentation of datasets with different acquisition protocols. The good quality of BOUNTI cortical segmentations also suggests that they can be used for automated surface-based analysis Makropoulos et al. (2018). However, any quantitative volumetry analysis of datasets from different studies will still require implementation of a dedicated harmonisation solution. The BOUNTI segmentation pipeline docker is publicly available online.

## Limitations and future work

In terms of the limitations, unlike the previously presented fetal brain atlas parcellation map in Gholipour et al. (2017), the current version of the BOUNTI pipeline includes only global tissue segmentation protocol with 19 ROIs without further subdivision into standard anatomical regions (e.g., frontal lobe). We are planning to further extend the parcellation map by separating the tissue compartments into anatomical ROIs and transient fetal compartments at the next stage of optimisation of the BOUNTI pipeline.

Any large scale application of the pipeline will also require retraining on a wider range of anomalies Fidon et al. (2021b) with potential optimisation of the parcellation protocol (e.g., for agenesis of the corpus callosum).

Moreover, while this work provides a baseline pipeline for further development, we did not perform any quantitative investigations of the impact of changes in intensity or image quality (i.e., definition of features) on segmentation results. Application of deep learning segmentation for multi-centre/scanner quantitative volumetry studies would require careful analysis of harmonisation requirements and potential optimisation of the networks.

Integration of data harmonisation Grigorescu et al. (2021) for different acquisition parameters and suboptimal image quality is one of the planned future steps along with incorporation of additional topological cortical constrains.

Further development of the BOUNTI pipeline would also require integration of automated image and segmentation quality control steps as well definition of what constitutes a correct segmentation and the acceptable levels of accuracy. This will also include a comprehensive quantitative evaluation of feasibility of using BOUNTI pipeline for analysis of different cohorts.

## 6. Conclusions

In this work, we formalised the new refined brain tissue protocol for 3D motion-corrected T2w fetal MRI in the spatiotemporal fetal atlas from the dHCP project. This protocol was used as a basis for training a deep learning BOUNTI pipeline for automated fetal brain segmentation on 360 fetal MRI datasets with different acquisition parameters. We used a semi-supervised approach for generation of the training datasets with manually refined labels propagated from the atlas. The pipeline showed robust performance across 21 - 36 weeks GA range and with different acquisition protocols.

The BOUNTI pipeline was then used to segment 390 normal control cases from 3 different cohorts with different acquisition parameters. Only minor errors in *<* 15% of cases thus significantly reducing the need for manual refinement. A preliminary analysis of the growth charts (21-38 weeks GA range) revealed no significant differences in the major solid tissue structures between the cohorts. In addition, comparison between 65 cases with ventriculomegaly and 60 normal control cases was in agreement with the findings reported in our earlier work using manual segmentationKyriakopoulou et al. (2014). These initial results suggest the general feasibility of using this pipeline as a basis for further development of fetal brain MRI parcellation tools and large-scale volumetric analysis.

The BOUNTI pipeline docker is publicly available online at SVRTK fetal segmentation repository^8^.

Our future work will focus on further extension of the anatomical parcellation map and optimisation of the BOUNTI pipeline for a wider range of brain anomalies as well as harmonisation, advanced cortex segmentation and segmentation quality control.

## Acknowledgements

We thank everyone who was involved in acquisition and analysis of the datasets at the Department of Perinatal Imaging and Health at Kings College London and St Thomas’ Hospital. We thank all participants and their families.

This work was supported by MRC Confidence in concept [MC PC 19041], the European Research Council under the European Union’s Seventh Framework Programme [FP7/ 20072013]/ERC grant agreement no. 319456 Dhcp project, the Wellcome Trust and EPSRC IEH award [102431] for the iFIND project, the NIH Human Placenta Project grant [1U01HD087202-01], MRC UK grant [MR/V002465/1], the Wellcome/ EPSRC Centre for Medical Engineering at King’s College London [WT 203148/Z/16/Z], Comunidad de Madrid-Spain under the line support for R&D projects for Beatriz Galindo researchers, MICIN / AEI / 10.13039/501100011033 / FEDER, EU, under Project PID2021-129022OA-I00, NIHR Advanced Fellowship awarded to Lisa Story [NIHR30166], the NIHR Clinical Research Facility (CRF) at Guy’s and St Thomas’ and by the National Institute for Health Research Biomedical Research Centre based at Guy’s and St Thomas’ NHS Foundation Trust and King’s College London.

The views expressed are those of the authors and not necessarily those of the NHS, the NIHR or the Department of Health.

## Author contributions

AUU prepared the datasets, developed the pipeline and prepared the manuscript; VK created the original parcellation protocol and contributed to development and testing of the pipeline; AM provided the initial dHCP segmentation pipeline; AG contributed to development the parcellation protocol and testing of the pipeline; DC contributed to training of the networks and testing of the pipeline; AD contributed to training of the networks; LCG contributed to preparation of the datasets; ANP contributed to acquisition of the datasets; IG contributed to development of the pipeline; LG contributed to analysis of the results; ER contributed to analysis of the results; DL provided fetal MRI datasets; KP provided fetal MRI datasets; SJC provided fetal MRI datasets; JH: provided fetal MRI datasets; LS contributed to testing of the pipeline and provided fetal MRI datasets; ADE provided fetal MRI datasets; MR contributed to conceptualisation of the project, development of the parcellation protocol and provided fetal MRI datasets; JVH provided fetal MRI datasets and contributed to analysis of the results; MD contributed to conceptualisation of the project and analysis of the results. All authors reviewed the manuscript.

## Footnotes

2 SVRTK docker (auto-2.00): https://hub.docker.com/r/fetalsvrtk/svrtk, https://github.com/SVRTK/SVRTK

3 ITK-SNAP tool: http://www.itksnap.org/

4 MIRTK toolbox: https://github.com/BioMedIA/MIRTK

5 dHCP fetal brain MRI atlas repository: https://gin.g-node.org/kcl_cdb/fetal_brain_mri_atlas

6 BOUNTI automated segmentation docker: https://hub.docker.com/r/fetalsvrtk/segmentation tag brain_bounti_tissue

7 dCHP fetal brain MRI atlas repository: https://gin.g-node.org/kcl_cdb/fetal_brain_mri_atlas

8 BOUNTI automated segmentation docker: https://hub.docker.com/r/fetalsvrtk/segmentation tag brain_bounti_tissue

## References

Aertsen, M., Diogo, M.C., Dymarkowski, S., Deprest, J., Prayer, D., 2020. Fetal mri for dummies: what the fetal medicine specialist should know about acquisitions and sequences. Prenatal Diagnosis 40, 6–17. doi:10.1002/pd.5579.

Bayer, S., Altman, J., 2003. The Human Brain During the Third Trimester. doi:10.1201/9780203494943.

Bayer, S., Altman, J., 2005. The Human Brain During the Second Trimester. doi:10.1201/9780203507483.

Cardoso, M.J., Li, W., Brown, R., Ma, N., Kerfoot, E., Wang, Y., Murrey, B., Myronenko, A., Zhao, C., Yang, D., et al., 2022. Monai: An open-source framework for deep learning in health-care. arXiv preprint arXiv:2211.02701 doi:10.48550/arXiv.2211.02701.

Cordero-Grande, L., et al., 2019. Automating Motion Compensation in 3T Fetal Brain Imaging: Localize, Align and Reconstruct, in: ISMRM 2019, p. 1000.

Dou, H., Karimi, D., Rollins, C.K., Ortinau, C.M., Vasung, L., Velasco-Annis, C., Ouaalam, A., Yang, X., Ni, D., Gholipour, A., 2021. A deep attentive convolutional neural network for automatic cortical plate segmentation in fetal mri. IEEE Transactions on Medical Imaging 40, 1123–1133. doi:10.1109/TMI.2020.3046579.

de Dumast, P., Kebiri, H., Atat, C., Dunet, V., Koob, M., Cuadra, M.B., 2021. Segmentation of the cortical plate in fetal brain mri with a topological loss, in: UNSURE and PIPPI MICCAI workshop 2021, Springer International Publishing. pp. 200–209. doi:10.1007/978-3-030-87735-4_19.

Ebner, M., Wang, G., Li, W., Aertsen, M., Patel, P.A., Aughwane, R., Melbourne, A., Doel, T., Dymarkowski, S., De Coppi, P., David, A.L., Deprest, J., Ourselin, S., Vercauteren, T., 2020. An automated framework for localization, segmentation and super-resolution reconstruction of fetal brain MRI. NeuroImage 206. doi:10.1016/j.neuroimage.2019.116324.

Fidon, L., Aertsen, M., Emam, D., Mufti, N., Guffens, F., Deprest, T., Demaerel, P., David, A.L., Melbourne, A., Ourselin, S., Deprest, J., Vercauteren, T., 2021a. Label-set loss functions for partial supervision: Application to fetal brain 3d mri parcellation, in: MICCAI 2021, Springer International Publishing. pp. 647–657. doi:10.1007/978-3-030-87196-3_60.

Fidon, L., Aertsen, M., Mufti, N., Deprest, T., Emam, D., Guffens, F., Schwartz, E., Ebner, M., Prayer, D., Kasprian, G., David, A.L., Melbourne, A., Ourselin, S., Deprest, J., Langs, G., Vercauteren, T., 2021b. Distributionally robust segmentation of abnormal fetal brain 3d mri, in: MICCAIN UNSURE, PIPPI work-shops 2021, pp. 263–273. doi:10.1007/978-3-030-87735-4_25.

Gholipour, A., Estroff, J.A., Warfield, S.K., 2010. Robust super-resolution volume reconstruction from slice acquisitions: Application to fetal brain mri. IEEE Transactions on Medical Imaging 29, 1739–1758. doi:10.1109/TMI.2010.2051680.

Gholipour, A., Rollins, C.K., Velasco-Annis, C., Ouaalam, A., Akhondi-Asl, A., Afacan, O., Ortinau, C.M., Clancy, S., Limperopoulos, C., Yang, E., Estroff, J.A., Warfield, S.K., 2017. A normative spatiotemporal mri atlas of the fetal brain for automatic segmentation and analysis of early brain growth. Nature: Scientific Reports 7, 1–13. doi:10.1038/s41598-017-00525-w.

Grigorescu, I., Vanes, L., Uus, A., Batalle, D., Cordero-Grande, L., Nosarti, C., Edwards, A.D., Hajnal, J.V., Modat, M., Deprez, M., 2021. Harmonized segmentation of neonatal brain mri. Frontiers in Neuroscience 15, 565. doi:10.3389/fnins.2021.662005.

Isensee, F., Jaeger, P.F., Kohl, S.A.A., Petersen, J., Maier-Hein, K.H., 2021. nnu-net: a self-configuring method for deep learning-based biomedical image segmentation. Nature Methods 18, 203–211. doi:10.1038/s41592-020-01008-z.

Karimi, D., Rollins, C.K., Velasco-Annis, C., Ouaalam, A., Gholipour, A., 2023. Learning to segment fetal brain tissue from noisy annotations. Medical Image Analysis, 102731 doi:10.1016/j.media.2022.102731.

Khalili, N., Lessmann, N., Turk, E., Claessens, N., de Heus, R., Kolk, T., Viergever, M.A., Benders, M., Iasgum, I., 2019. Automatic brain tissue segmentation in fetal mri using convolutional neural networks. Magnetic Resonance Imaging 64, 77–89. doi:10.1016/j.mri.2019.05.020.

Kuklisova-Murgasova, M., Quaghebeur, G., Rutherford, M.A., Hajnal, J.V., Schnabel, J.A., 2012. Reconstruction of fetal brain mri with intensity matching and complete outlier removal. Med Image Analysis 16, 1550–1564. doi:10.1016/j.media.2012.07.004.

Kyriakopoulou, V., Vatansever, D., Davidson, A., Patkee, P., Elkommos, S., Chew, A., Martinez-Biarge, M., Hagberg, B., Damodaram, M., Allsop, J., Fox, M., Hajnal, J.V., Rutherford, M.A., 2017. Normative biometry of the fetal brain using magnetic resonance imaging. Brain Structure and Function 222, 2295–2307. doi:10.1007/s00429-016-1342-6.

Kyriakopoulou, V., Vatansever, D., Elkommos, S., Dawson, S., McGuinness, A., Allsop, J., Molnér, Z., Hajnal, J., Rutherford, M., 2014. Cortical overgrowth in fetuses with isolated ventriculomegaly. Cerebral Cortex 24, 2141–2150. doi:10.1093/cercor/bht062.

Li, L., Ma, Q., Li, Z., Ouyang, C., Zhang, W., Price, A., Kyriakopoulou, V., Grande, L.C., Makropoulos, A., Hajnal, J., Rueckert, D., Kainz, B., Alansary, A., 2022. Fetal cortex segmentation with topology and thickness loss constraints, in: MICCAI EPIMI, ML-CDS, TDA4BioMedicalImaging workshop 2022, pp. 123–133. doi:10.1007/978-3-031-23223-7_11.

Li, L., Sinclair, M., Makropoulos, A., Hajnal, J.V., David Edwards, A., Kainz, B., Rueckert, D., Alansary Amir”, e.C.H., Licandro, R., Baumgartner, C., Melbourne, A., Dalca, A., Hutter, J., Tanno, R., Abaci Turk, E., Van Leemput, K., Torrents Barrena, J., Wells, W.M., Macgowan, C., 2021. Cas-net: Conditional atlas generation and brain segmentation for fetal mri, in: UNSURE and PIPPI MICCAI workshops, Springer Nature Switzerland. pp. 221–230. doi:10.1007/978-3-030-87735-4_21.

Machado-Rivas, F., Gandhi, J., Choi, J.J., Velasco-Annis, C., Afacan, O., Warfield, S.K., Gholipour, A., Jaimes, C., 2021. Normal growth, sexual dimorphism, and lateral asymmetries at fetal brain mri. Radiology, 211222 doi:10.1148/radiol.211222.

Makropoulos, A., Robinson, E.C., Schuh, A., Wright, R., Fitzgibbon, S., Bozek, J., Counsell, S.J., Steinweg, J., Vecchiato, K., Passerat-Palmbach, J., Lenz, G., Mortari, F., Tenev, T., Duff, E.P., Bastiani, M., Cordero-Grande, L., Hughes, E., Tusor, N., Tournier, J.D., Hutter, J., Price, A.N., Teixeira, R.A.P.G., Murgasova, M., Victor, S., Kelly, C., Rutherford, M.A., Smith, S.M., Edwards, A.D., Hajnal, J.V., Jenkinson, M., Rueckert, D., 2018. The developing human connectome project: a minimal processing pipeline for neonatal cortical surface reconstruction. Neuroimage 173, 88–112. doi:10.1101/125526.

Oktay, O., Schlemper, J., Folgoc, L.L., Lee, M., Heinrich, M., Misawa, K., Mori, K., Mcdonagh, S., Hammerla, N.Y., Kainz, B., Glocker, B., Rueckert, D., 2018. Attention u-net: Learning where to look for the pancreas, in: MIDDL 2016.

Payette, K., de Dumast, P., Kebiri, H., Ezhov, I., Paetzold, J.C., Shit, S., Iqbal, A., Khan, R., Kottke, R., Grehten, P., Ji, H., Lanczi, L., Nagy, M., Beresova, M., Nguyen, T.D., Natalucci, G., Karayannis, T., Menze, B., Cuadra, M.B., Jakab, A., 2021. An automatic multi-tissue human fetal brain segmentation benchmark using the fetal tissue annotation dataset. Scientific Data 8, 1–14. doi:10.1038/s41597-021-00946-3.

Payette, K., Kottke, R., Jakab, A., 2020. Efficient multi-class fetal brain segmentation in high resolution mri reconstructions with noisy labels, in: MICCAI PIPPI workshop 2020, pp. 295–304. doi:10.1007/978-3-030-60334-2_29.

Payette, K., Li, H., de Dumast, P., Licandro, R., Ji, H., Siddiquee, M.M.R., Xu, D., Myronenko, A., Liu, H., Pei, Y., Wang, L., Peng, Y., Xie, J., Zhang, H., Dong, G., Fu, H., Wang, G., Rieu, Z., Kim, D., Kim, H.G., Karimi, D., Gholipour, A., Torres, H.R., Oliveira, B., Vilaça, J.L., Lin, Y., Avisdris, N., Ben-Zvi, O., Bashat, D.B., Fidon, L., Aertsen, M., Vercauteren, T., Sobotka, D., Langs, G., Alenyà, M., Villanueva, M.I., Camara, O., Fadida, B.S., Joskowicz, L., Weibin, L., Yi, L., Xuesong, L., Mazher, M., Qayyum, A., Puig, D., Kebiri, H., Zhang, Z., Xu, X., Wu, D., Liao, K., Wu, Y., Chen, J., Xu, Y., Zhao, L., Vasung, L., Menze, B., Cuadra, M.B., Jakab, A., 2022. Fetal brain tissue annotation and segmentation challenge results. doi:10.48550/arXiv.2204.09573.

Pei, Y., Chen, L., Zhao, F., Wu, Z., Zhong, T., Wang, Y., Chen, C., Wang, L., Zhang, H., Wang, L., Li, G., 2021. Learning spatiotemporal probabilistic atlas of fetal brains with anatomically constrained registration network, in: MICCAI 2021, pp. 239–248. doi:10.1007/978-3-030-87234-2_23.

Prayer, D., Malinger, G., Brugger, P.C., Cassady, C., Catte, L.D., Keersmaecker, B.D., Fernandes, G.L., Glanc, P., Gonçalves, L.F., Gruber, G.M., Laifer-Narin, S., Lee, W., Millischer, A.E., Molho, M., Neelavalli, J., Platt, L., Pugash, D., Ramaekers, P., Salomon, L.J., Sanz, M., Timor-Tritsch, I.E., Tutschek, B., Twickler, D., Weber, M., Ximenes, R., Raine-Fenning, N., 2017. Isuog practice guidelines: performance of fetal magnetic resonance imaging. Ultrasound in Obstetrics and Gynecology 49, 671–680. doi:10.1002/uog.17412.

Price, A.N., Cordero-grande, L., Hughes, E., Hiscocks, S., Green, E., Mccabe, L., Ferrazzi, G., Deprez, M., Roberts, T., Christiaens, D., Duff, E., Karolis, V., Malik, J., Rutherford, M.A., Edwards, D.A., Hajnal, J.V., 2019. The developing human connectome project (dhcp): fetal acquisition protocol, in: ISMRM 2019.

Rollins, C.K., Ortinau, C.M., Stopp, C., Friedman, K.G., Tworetzky, W., Gagoski, B., Velasco-Annis, C., Afacan, O., Vasung, L., Beaute, J.I., Rofeberg, V., Estroff, J.A., Grant, P.E., Soul, J.S., Yang, E., Wypij, D., Gholipour, A., Warfield, S.K., Newburger, J.W., 2021. Regional brain growth trajectories in fetuses with congenital heart disease. Annals of Neurology 89, 143–157. doi:10.1002/ana.25940.

Ronneberger, O., Fischer, P., Brox, T., 2015. U-net: Convolutional networks for biomedical image segmentation, in: MICCAI 2015, pp. 234–241.

Royston, P., Wright, E., 1998. How to construct ‘normal ranges’ for fetal variables. Ultrasound in Obstetrics Gynecology 11, 30–38. doi:10.1046/j.1469-0705.1998.11010030.x.

Rueckert, D., Sonoda, L.I., Hayes, C., Hill, D.L.G., Leach, M.O., Hawkes, D.J., 1999. Nonrigid registration using free-form deformations: Application to breast mr images. IEEE TRANSACTIONS ON MEDICAL IMAGING 18, 712–721. doi:10.1109/42.796284.

Rutherford, M., Jiang, S., Allsop, J., Perkins, L., Srinivasan, L., Hayat, T., Kumar, S., Hajnal, J., 2008. Mr imaging methods for assessing fetal brain development. Developmental Neurobiology 68, 700–711. doi:10.1002/dneu.20614.

Story, L., Davidson, A., Patkee, P., Fleiss, B., Kyriakopoulou, V., Colford, K., Sankaran, S., Seed, P., Jones, A., Hutter, J., Shennan, A., Rutherford, M., 2021. Brain volumetry in fetuses that deliver very preterm: An mri pilot study. NeuroImage: Clinical 30, 102650. doi:10.1016/j.nicl.2021.102650.

Uus, A., Kyriakopoulou, V., Cordero Grande, L., Christiaens, D., Pietsch, M., Price, A., Wilson, S., Patkee, P., Karolis, S., Schuh, A., Gartner, A., Williams, L., Hughes, E., Arichi, T., O’Muircheartaigh, J., Hutter, J., Robinson, E., Tournier, J., Rueckert, D., Counsell, S., Rutherford, M., Deprez, M., Hajnal, J., Edwards, A., 2023. Multi-channel spatio-temporal mri atlas of the normal fetal brain development from the developing human connectome project. G-Node doi:10.12751/g-node.ysgsy1.

Uus, A.U., Collado, A.E., Roberts, T.A., Hajnal, J.V., Rutherford, M.A., Deprez, M., 2022. Retrospective motion correction in foetal mri for clinical applications: existing methods, applications and integration into clinical practice. The British Journal of Radiology doi:10.1259/bjr.20220071.

Vasung, L., Rollins, C.K., Zhang, J., Velasco-Annis, C., Yang, E., Lin, P.Y., Sutin, J., Warfield, S.K., Soul, J., Estroff, J., Connolly, S., Barnewolt, C., Gholipour, A., Feldman, H.A., Grant, P.E., 2022. Abnormal development of transient fetal zones in mild isolated fetal ventriculomegaly. Cerebral Cortex, bhac125 doi:10.1093/cercor/bhac125.

Wright, R., Khanal, B., Gomez, A., Skelton, E., Matthew, J., Hajnal, J.V., Rueckert, D., Schnabel, J.A., 2018. Lstm spatial cotransformer networks for registration of 3d fetal us and mr brain images, in: MICCAI 2018, Springer International Publishing. pp. 107–116. doi:10.1007/978-3-030-00807-9.

Yushkevich, P.A., Piven, J., Hazlett, H.C., Smith, R.G., Ho, S., Gee, J.C., Gerig, G., 2006. User-guided 3d active contour segmentation of anatomical structures: Significantly improved efficiency and reliability. NeuroImage 31, 1116–1128. doi:10.1016/j.neuroimage.2006.01.015.

Özgün Ç içek, Abdulkadir, A., Lienkamp, S.S., Brox, T., Ronneberger, O., 2016. 3d u-net: Learning dense volumetric segmentation from sparse annotation, in: MICCAI 2016, pp. 424–432. doi:10.1007/978-3-319-46723-8_49.

